# Monomeric streptavidin: a versatile regenerative handle for force spectroscopy

**DOI:** 10.1101/276444

**Authors:** Magnus S. Bauer, Lukas F. Milles, Steffen M. Sedlak, Hermann E. Gaub

## Abstract

Most avidin-based handles in force spectroscopy are tetravalent biotin binders. Tetravalency presents two issues: multiple pulling geometries as well as multiple targets bound simultaneously. Additionally, such tetravalent handles require elaborate purification protocols in order to reassemble. A stoichiometric, monomeric variant of streptavidin (mcSA2) had been engineered previously. It is readily expressed and purified, and it binds biotin with a nanomolar K_D_. For atomic force microscopy-based single-molecule force spectroscopy (AFM-SMFS), we fused the monomeric streptavidin with a small protein domain as an experimental fingerprint and to improve solubility. A ybbR-tag was additionally included for covalent site-specific tethering. Rupture forces of the mcSA2:biotin complex were found to be in a comparable range above 150 pN at force loading rates of 1E4 pN/s as for previously published, tetravalent streptavidin:biotin systems. Additionally, when tethering mcSA2 from its C-terminus, rupture forces were found to be slightly higher than when tethered N-terminally. Due to its monomeric nature, mcSA2 could also be chemically denatured and subsequently refolded - and thus regenerated during an experiment, in case the handle gets misfolded or clogged. We show that mcSA2 features a straightforward expression and purification with flexible tags, high stability, regeneration possibilities and an unambiguous pulling geometry. Combined, these properties establish mcSA2 as a reliable handle for single-molecule force spectroscopy.

## Introduction

Avidin-based handles have a long and successful history in biotechnology. They are widely applied as tagging and pull-down handles due to their femtomolar affinity towards the small molecule biotin, low off-rate, broad availability, and easy handling. As the first receptor-ligand system probed in atomic force microscopy-based single-molecule force spectroscopy (AFM-SMFS) studies (1,2), they still enjoy great popularity as handles to apply force to biomolecular systems.

Avidin (3) and similar molecules, such as streptavidin (4) or strep-tactin (5), are tetramers composed of four separate subunits, each capable of binding a single biotin molecule with high affinity. However, for some applications there is yet a need for precise control over stoichiometry. Considerable effort went into the design of a monovalent variant of streptavidin, a tetramer with only one single biotin binding subunit (6). For SMFS studies, an identical approach guaranteeing a well-defined tethering with 1:1 binding stoichiometry and specific pulling geometry was pursued by assembling a functional streptavidin subunit with three non-functional subunits (7). An analogous approach has been established for strep-tactin to tether a single strep-tag II peptide (8). These approaches achieve monovalent binding behavior but still require tetrameric structure to retain function. Additionally, they rely on elaborate purification procedures to assemble the tetrameric structure.

Recently, Park and colleagues undertook the effort to engineer a monomeric streptavidin - a solitary, yet functional streptavidin subunit. Monomeric variants inherently have some disadvantages compared to their tetrameric equivalents, among them lower biotin affinity, low solubility and problems with aggregation (9,10). To overcome these issues, Lim et al. engineered a monomeric streptavidin (mcSA) as a chimera based on structural homology modeling of streptavidin and rhizavidin, a dimeric protein that binds biotin using residues from only a single subunit (11). The resulting biotin affinity of 2.8 nM is the highest among non-tetrameric streptavidin. DeMonte et al. crystalized mcSA, analyzed it in detail, and improved it further by some mutations in the binding pocket (12). The resulting mcSA2 has a 20-40% lower off-rate. Adding solubility tags optimized the expression procedure (13).

In this study, we employ mcSA2 and combine it with the 4^th^ filamin domain from *Dictyostelium discoideum* (ddFLN4) as both a molecular fingerprint for SMFS and a solubility enhancer. Additionally, an N- or C-terminal polyhistidine purification tag and a ybbR-tag (14) for site-specific covalent immobilization were included. We describe a straightforward expression and purification protocol under denaturing conditions to eliminate biotin already present in the binding pocket beforehand, followed by refolding of the fusion protein via dialysis. We test the new mcSA2 force handle in AFM-SMFS and show that the mcSA2:biotin complex withstands forces comparable to the streptavidin:biotin interaction and is also showing two different force regimes by pulling from the molecule’s N- or C-terminus. Additionally, the monomeric nature of the employed handles entail a unique feature: it can be completely denatured and refolded *in situ* making it superior to tetrameric biotin handles. For example, if clogged by stray biotin or trapped in misfolded states, the mcSA2 handle can be regenerated by recovering its binding ability. This property results in higher data yield and better statistics as it allows performing AFM-SMFS experiments with a single cantilever for several days without loss of interaction.

## Results and Discussion

### Applicability of the handle for force spectroscopy

To probe the applicability and long term stability of mcSA2 as a handle for force spectroscopy AFM-SMFS measurements were performed. We investigated two similar constructs to examine the mechanical characteristics of the unbinding of biotin from mcSA2 under force application on its different termini: an mcSA2 with the ddFLN4 fingerprint and the ybbR-tag on the N-terminus (geometry N, ybbR-ddFLN4-mcSA2) and an mcSA2 with the fingerprint domain and the immobilization tag on its C-terminus (geometry C, mcSA2-ddFLN4-ybbR) as depicted in Figure 1A,B.

**Figure 1.**
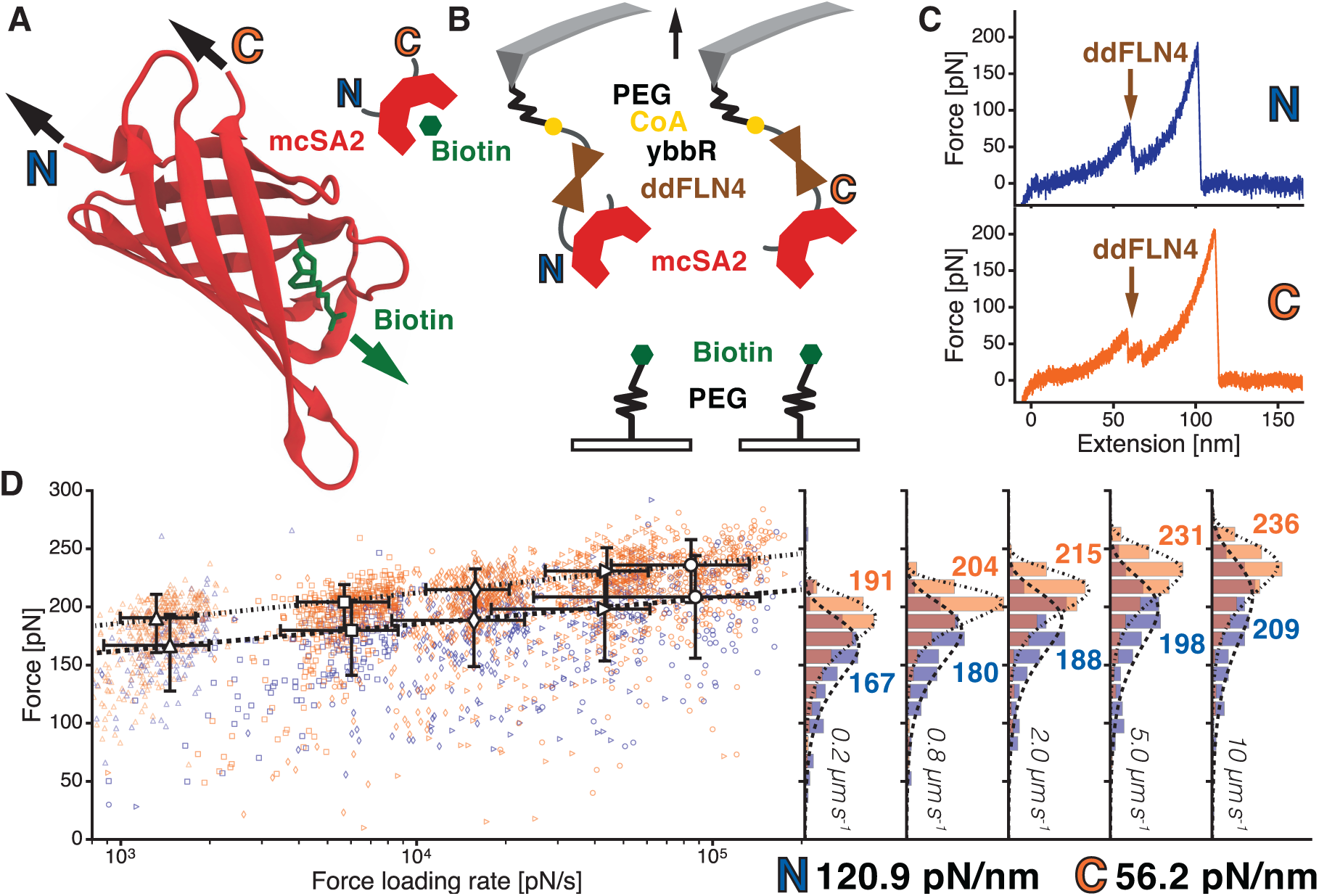
Characterization of the mcSA2 handle by AFM-based SMFS. Panel A: the crystal structure adapted from protein database (PDB) entry 4JNJ (12) and schematic of mcSA2 (red) and biotin (green) with pulling geometries N (blue, pulled from N-terminus) and C (orange, pulled from C-terminus). Panel B: a schematic of the attachment chemistry is depicted. Both constructs are immobilized on an aminosilanized cantilever with heterobifunctional NHS-PEG-maleimide linkers. On the maleimide side of the PEG, a CoA is attached for an sfp phosphopantetheinyl transferase (sfp) catalyzed reaction with the ybbR-tag of the mcSA2 handle constructs. The likewise aminosilanzed glass surface is functionalized with a heterobifunctional NHS-PEG-biotin linker. Panel C: two exemplary curves for both geometries N (top) and C (bottom) with its characteristic ddFLN4 fingerprint. Panel D: a dynamic force spectrum and force histograms of both geometries N (blue) and C (orange) indicating a similar force loading rate dependence but with generally higher forces for geometry C. The forces indicated in the histograms show the most probable force in pN according to the Bell-Evans-model. In this experimental setup the different force datasets had to be recorded with two separate cantilevers in order to probe the long term stability of the handles in both geometries on the cantilever. Since e.g. deviations in the cantilevers’ spring constants (bottom right) hinder to compare forces directly in absolute values, both tethering geometries were additionally measured with a single cantilever in one measurement for better comparability as shown in Figure 2.

The handles were covalently linked to AFM cantilevers and probed against a biotinylated surface (cf. materials and methods, Figure 1B). Single unbinding events could be identified by the characteristic unfolding pattern of ddFLN4, which includes a shielded substep (Figure 1C). The recurring unfolding pattern assured that the large number of specific mcSA2:biotin interaction events are pulled specifically by a single handle in a well-defined geometry, and thus shows that the handle can be implemented as a reliable force handle in SMFS experiments. The resulting forces of 150-200 pN needed for detaching a single biotin from the mcSA2 binding pocket are comparable to what has been reported for the streptavidin:biotin interaction (1,7,15). Using different retraction velocities, a dynamic force spectrum was obtained and fitted as a single bond dissociation over an energy barrier according to Bell (16) and Evans (17). For geometry N, the fit yielded a distance to the transition state x_0_ = 0.42 nm and a zero-force off-rate k_off,0_ = 7.7 × 10^-6^ s^-1^. For geometry C, x_0_ = 0.37 nm and k_off,0_ = 6.1 × 10^-6^ s^-1^ were obtained. Over the broad range of loading rates, unbinding forces for the C-terminally tethered mcSA2 are higher than those for the N-terminally tethered mcSA2 as correctly as it could be determined with two different cantilevers.

### Comparison of N- and C-terminal pulling geometry

Calibration errors and changes in force due to differing spring constants between individual cantilevers can render comparison of experimental force data – especially when addressing small force differences – unreliable. To compare rupture forces of mcSA2:biotin loaded in geometry N and C, we thus performed measurements with one single cantilever by immobilizing the two different constructs of the mcSA2 handle at two separate spots on one functionalized glass slide (Figure 2A). This way both geometries can be probed with the same cantilever with one consistent spring constant of 139.2 pN/nm in order to yield directly comparable force values. To ensure single-molecule interactions, we introduced an additional fingerprint domain on the cantilever: the refolding, alpha-helical protein FIVAR (derived from “Found In Various Architectures”) domain (18) from the pathogen C*lostridium perfringens* that is known to unfold at forces of 50-60 pN (Figure 2B). Biotinylation was accomplished using an AviTag sequence (19), which is covalently modified with a biotin during protein expression (cf. Materials and Methods). Covalent and site-specific tethering was again achieved employing a ybbR-tag.

**Figure 2.**
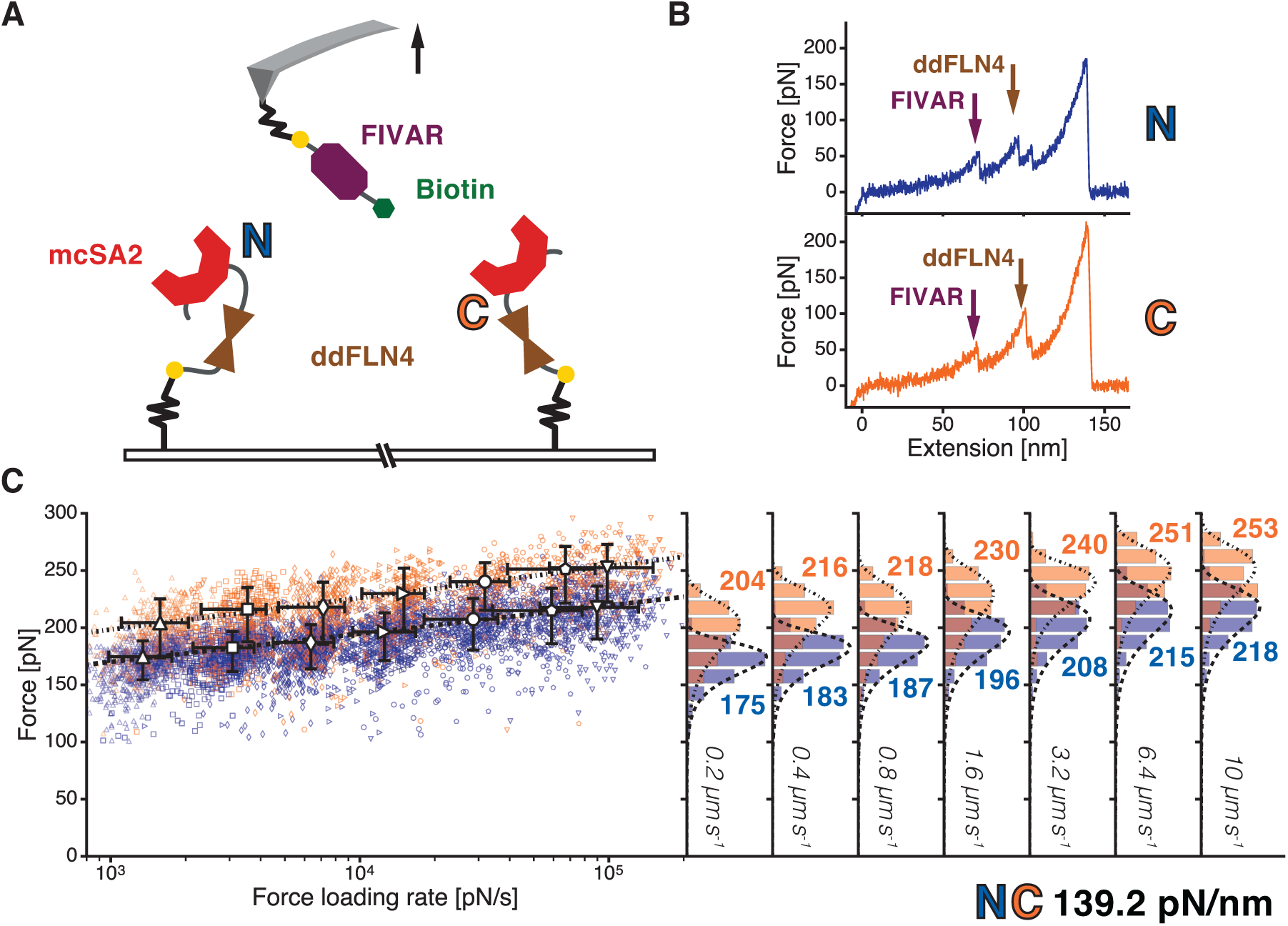
Direct comparison of unbinding forces for two different tethering geometries N and C. Panel A: to compare the unbinding forces of the two tethering scenarios, both geometries N (blue, pulled from N-terminus) and C (orange, pulled from C-terminus) were immobilized on separate spots on a surface and were probed using the same cantilever harboring a FIVAR domain with a Biotin attached. Panel B: two exemplary curves for both geometries N (top) and C (bottom) with its characteristic FIVAR and ddFLN4 fingerprint. Panel C: the data were recorded within one experiment by switching between the two spots every 300 curves. This resulted in a dynamic force spectrum and force histograms for both geometries, allowing direct comparison of unbinding forces for both geometries N and C. The forces indicated in the histograms show the most probable force in pN according to the Bell-Evans-model. The spring constant of the cantilever (139.2 pN/nm) used to pull both geometries is shown on the bottom right.

In this SMFS experiment, the cantilever alternated between surface areas with mcSA2 tethered in geometry N and C for every 300 approaches. While the unfolding forces of the fingerprint domains remained the same for both tethering geometries, we found the mcSA2:biotin interaction to be significantly stronger for geometry C than for geometry N throughout all varied retraction velocities. The most probable rupture forces in pN according to the Bell-Evans-model for each geometry is shown in Figure 2C. The most probable forces for geometry C consistently exceeded those for geometry N by 30 – 40 pN. Fitting the dynamic force spectrum with the Bell-Evans-model, the N-terminal tethering yielded a distance to the transition state x_0_ = 0.39 nm and a zero-force off-rate k_off,0_ = 1.2 × 10^-5^ s^-1^, while x_0_ = 0.35 nm and k_off,0_ = 5.3 × 10^-6^ s^-1^ was obtained for the C-terminal tethering. These results agree well with the results obtained for the mcSA2 handles on the cantilever from Figure 1D.

### Characterization of affinity

To determine whether the difference in unbinding forces for the two different geometries emerges from the way the mcSA2 molecule is loaded or by a conformational difference resulting from the addition of ddFLN4 to the termini, we performed fluorescence anisotropy experiments. In a competition assay, we measured the off-rates for both constructs in solution, thus in the absence of external force (Figure 3). Measurements of mcSA2 with ddFLN4 on the N- and C-terminus yielded off-rates of 1.05 × 10^-4^ s^-1^ and 1.08 × 10^-4^ s^-1^, respectively. Regarding the measurement’s accuracy, the off-rates of both constructs are considered to be equal. Therefore, we conclude that the difference in unbinding force during AFM-SMFS is determined solely by the way force is applied to the handle and thus the trajectory chosen to overcome the binding energy barrier rather than the position of the ddFLN4 fingerprint itself.

**Figure 3.**
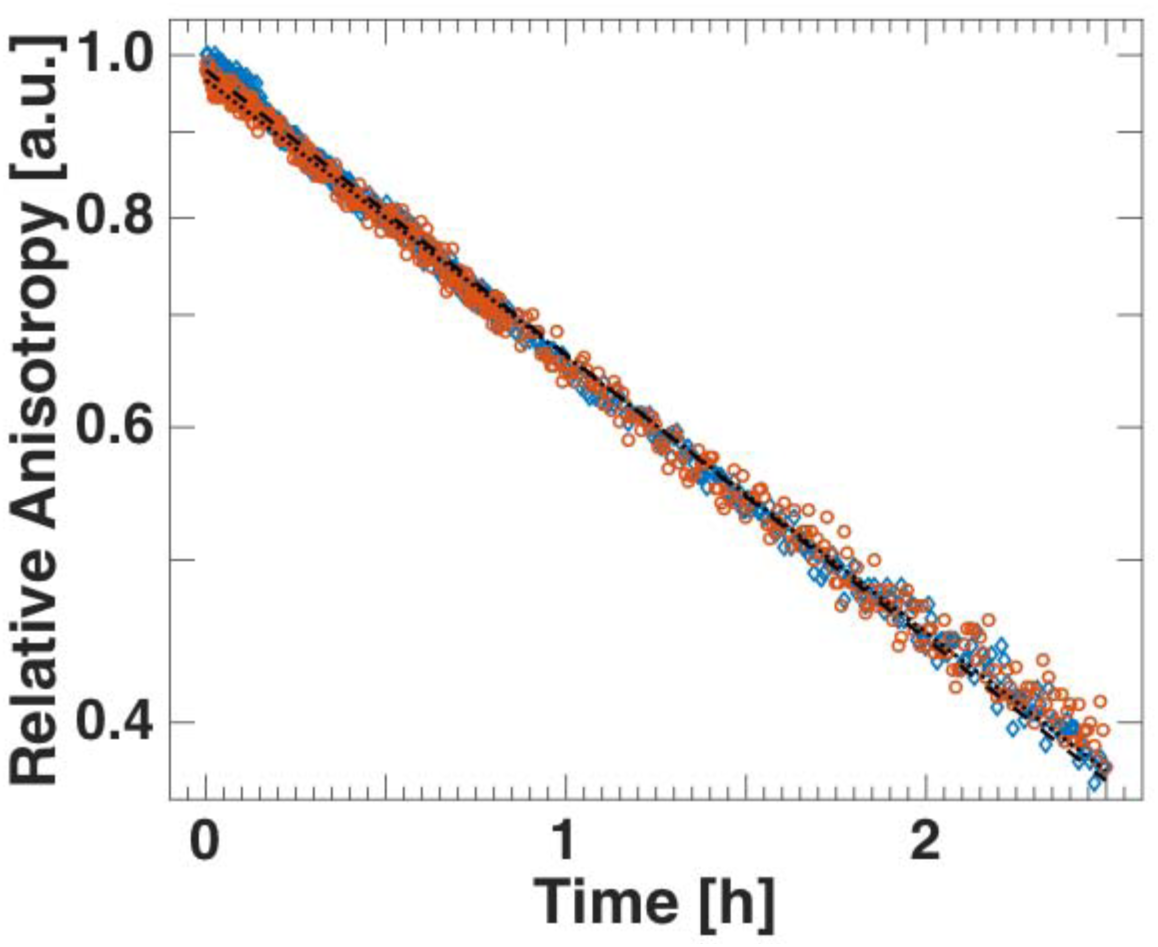
Off-rates for two different tethering geometries. For geometry C (orange circles) and geometry N (blue diamonds), the relative anisotropy is plotted over time. Fitting the off-rates yields 0.000108 s-1 × t - 0.208 for geometry C (black dotted line) and 0.000105 s-1 × t - 0.342 for geometry N (black dashed line). Hence, no significant difference for the off-rates is observed. (Here, relative anisotropy denotes the logarithm of the present anisotropy difference between sample and reference divided by the difference at the moment of biotin addition, t=0.)

### Regeneration of the mcSA2 handle

In AFM-SMFS experiments, a streptavidin handle on the cantilever may occasionally pick up biotinylated molecules that were unspecifically adsorbed to the sample surface. The high affinity of the streptavidin:biotin interaction is in this case particularly disadvantageous, because biotinylated molecules block the binding pockets of the handle almost irreversibly. Once a cantilever is clogged, the interaction with the biotinylated molecules on the surface is lost and they cannot be investigated any further. To regenerate such a clogged handle, we placed the cantilever in 6 M guanidine hydrochloride to denature the mcSA2 handle, releasing biotinylated molecules from its binding pocket. Subsequent gentle washing steps in phosphate buffered saline facilitates the refolding of the handle into its functional state. The ddFLN4 fingerprint also rapidly refolds. Using this protocol, we could recover mcSA2 from clogged or misfolded states and regain tethering activity on the surface.

In our experiment, we regenerated the handle up to 3 times but the regeneration steps are not limited to that. Resuming the SMFS measurement, no significant change in unfolding or rupture forces was detectable (Figure 4).

**Figure 4.**
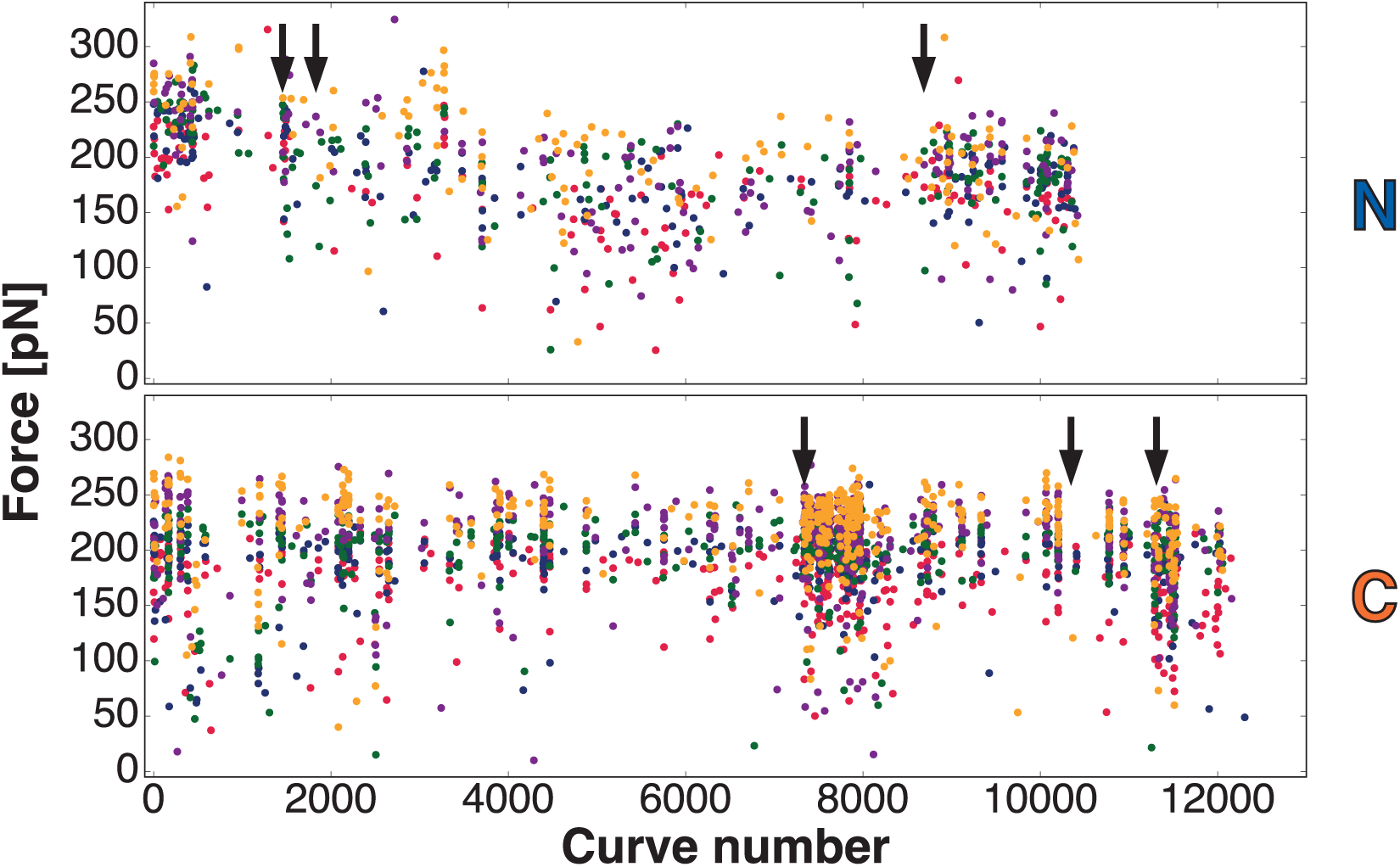
Regeneration of the mcSA2 handle. During the course of an AFM-SMFS measurement, the pulling handle eventually gets clogged with excess biotin picked up from the surface or is brought into a misfolded state rendering it unable to bind biotin any more. Due to its monomeric nature mcSA2 is able to be unfolded in 6 M guanidine hydrochloride and subsequently refolded in phosphate buffered saline in order to resume the measurement. These regeneration steps are indicated with black arrows. The Graph shows the force of mcSA2:biotin rupture in pN vs. curve number from the dataset shown in Figure 1D. Each curve number contains one pulling cycle of five retraction speeds of 200 nm/s (red), 800 nm/s (blue), 2000 nm/s (green), 5000 nm/s (purple), 10000 nm/s (orange). After a regeneration step, the ability to bind biotin is recovered - shown by the increased number of interactions recorded after the black arrows. This worked well with both geometries N (top panel) and C (bottom panel).

### Conclusion

Building on monomeric streptavidin, we could establish a highly specific handle for biotin-binding that is straightforward to produce and employ in force spectroscopy experiments. Additionally, mcSA2 is a long-lived tethering handle, enhanced in its performance even further as it can be regenerated by refolding. Our study shows that mcSA2 can be a significant asset for SMFS and related applications. Combined with site-specific anchoring, it permits high data yields, whenever biotinylation is possible.

We could also show the importance of anchoring positions for the stability of a receptor-ligand interaction since this changes the trajectory chosen in the binding energy landscape to overcome the energy barrier. Therefore precise control of the pulling geometry changes the interaction’s mechanostability, permitting to switch the addressed force range. In conclusion, its robustness and versatility renders mcSA2 an excellent choice for force spectroscopy measurements.

## Materials and Methods

### Protein Expression and Purification - Gene construction and cloning

mcSA2 was expressed and purified with a fingerprint and solubility enhancer, the 4^th^ filamin domain of *Dictyostelium discoideum* (ddFLN4). This small Ig-like fold expresses well and refolds rapidly. By varying the position of the ybbR-tag, used for covalent protein pulldown, two different tethering geometries could be examined: Geometry N with mcSA2 on the C-terminus (ybbR-ddFLN4-mcSA2) and geometry C with mcSA2 on the N-terminus (mcSA2-ddFLN4-ybbR). These constructs were cloned using the Gibson assembly strategy into pET28a vectors. The ybbR-HIS-FIVAR-AviTag was cloned into a pAC4 vector.

Both constructs were expressed in NiCo Cells (New England Biolabs) in autoinduction Media under Kanamycin resistance. Harvested cell pellets were resuspended in 50 mM TRIS, 50 mM NaCl, 10 % (w/v) Glycerol, 0.1 % (v/v) Triton X-100, 5 mM MgCl2 at pH 8.0. To enhance cell lysis, 100 µg/ml lysozyme and 10 µg/ml DNase were added. The solution was then sonicated for 2 x 8 min. The lysed cells were spun down for 10 min at 7000 rpm in a precooled centrifuge at 4°C. Solid guanidine hydrochloride was added to the supernatant to a concentration of 6 M to completely unfold the construct to release any bound biotin. The denatured construct was purified by immobilized metal ion affinity chromatography using a HisTrap FF column (GE Healthcare). Once the protein was bound to the column, it was extensively washed with denaturing buffer to remove any stray biotin present. Finally the protein was eluted with 200 mM Imidazole. The purified protein was refolded by three rounds of dialyzation against Phosphate buffered saline (PBS) overnight and finally, after the addition of 10% glycerol, flash frozen in liquid nitrogen, to be stored at -80°C.

ybbR-FIVAR-AviTag on a pAC4 vector was expressed in E. Coli CVB101 (Avidity LLC), supplemented with biotin in the expression medium in autoinduction media and was purified identically, although non-denaturing conditions.

### Surface functionalization for the AFM measurement

The preparation of the experiments comprises two similar immobilization protocols. Either for the mcSA2 or FIVAR-Biotin construct with ybbR-tag or the NHS-PEG-Biotin on a glass/cantilever surface. The experiments were designed to either have mcSA2 on the cantilever and NHS-PEG-Biotin or FIVAR-Biotin on the surface or vice versa. Immobilization of mcSA2 to cantilever or glass surface is identical to the protocol used for the attachment of FIVAR. (14,20)

### Preparation of Cantilevers

For aminosilanization of the cantilevers (BioLever Mini obtained from Olympus, Japan) they were first oxidized in a UV-ozone cleaner (UVOH 150 LAB, FHR Anlagenbau GmbH, Ottendorf-Okrilla, Germany) and subsequently silanized for 2 minutes in (3-Aminopropyl)dimethylethoxysilane (ABCR, Karlsruhe, Germany; 50 % v/v in Ethanol). For rinsing, the cantilevers were stirred in 2-Propanol (IPA), ddH_2_O and afterwards dried at 80°C for 30 minutes. After that the cantilevers were incubated in a solution of 25 mM heterobifunctional PEG spacer (MW 5000, Rapp Polymere, Tübingen, Germany) solved in 50 mM HEPES for 30 minutes. Subsequent to rinsing with ddH_2_O, the surfaces were incubated in 20 mM Coenzyme A (Calbiochem) dissolved in coupling buffer (sodium phosphate, pH 7.2) to react with the maleimide groups. After that the levers get rinsed with ddH_2_O. Then the ybbR-tag of the mcSA2 (at 5-50 µM) construct (in PBS supplemented with 10 mM MgCl_2_) is attached covalently by a sfp (at 2 µM) catalyzed reaction to the CoA. After 30 min to 2 h the protein is covalently connected resulting in an unambiguous, site-specific pulldown. Finally, the cantilevers were rinsed thoroughly and stored in 1 x PBS.

For the preparation of PEG Biotin (5000 Da) cantilevers pegylation protocols were identical, only that NHS-PEG-Biotin instead of NHS-PEG-Maleimide was applied for 1 h.

For the preparation of FIVAR cantilevers the mcSA2 construct was substituted for the FIVAR construct. Similar concentrations of protein were used.

### Preparation of Glass Surfaces

Before aminosilanization the glass surfaces were cleaned by sonication in 50 % (v/v) Isopropanol (IPA) in ultrapure water for 15 minutes. For oxidation the glass surfaces were soaked for 30 minutes in a solution of 50 % (v/v) hydrogen peroxide (30 %) and sulfuric acid. Afterwards they were thoroughly washed in ultrapure water and then blown dry in a gentle nitrogen stream. Silanization is achieved by incubating in (3-Aminopropyl)dimethylethoxysilane (ABCR, Karlsruhe, Germany, 1.8 % v/v in Ethanol) while gently shaking. Thereafter, surfaces were washed again in IPA and ultrapure water and then dried at 80°C for 40 minutes, to be stored under Argon for weeks.

To attach mcSA2 to the glass surface heterobifunctional Polyethyleneglycol (PEG, 5000 Da, dissolved in 100 mM HEPES pH 7.5 at 25 mM for 30 min) spacers were used to avoid unspecific interactions between the cantilever and the glass surface. The PEG spacers had an N-hydroxysuccinimide (NHS) group on one side, for attachment to the aminosilanized surface. The other end provided a Maleimide group for subsequent coupling to the thiol group of Coenzyme A (CoA, 1 mM in 50 mM sodium phospahte, 50 mM NaCl, 10 mM EDTA, pH 7.2, incubated for 1 h). Through a reaction catalyzed by sfp (at 2 µM) the CoA was covalently connected to the ybbR-tag of the mcSA2 (at 5-50 µM) construct (in PBS supplemented with 10 mM MgCl_2_ for 30 min to 2 h), resulting in an unambiguous, site-specific pulldown.

For the preparation of PEG Biotin (5000 Da) surfaces pegylation protocols were identical, only that NHS-PEG-Biotin instead of NHS-PEG-Maleimide was applied for 1 h.

For the preparation of FIVAR surfaces the mcSA2 construct was substituted for the FIVAR construct. Similar concentrations of protein were used.

### AFM-SMFS

Adapted from Milles et al. (18):

AFM-SMFS data was acquired on a custom-built AFM operated in closed loop by a MFP3D controller (Asylum Research, Santa Barbara, CA, USA) programmed in Igor Pro 6 (Wavemetrics, OR, USA). Cantilevers were briefly (<200 ms) and gently (< 200 pN) brought in contact with the functionalized surface and then retracted at constant velocities ranging from 0.2, 0.8, 1.6, 2.0, 3.2, 5.0, 6.4 to 10.0 µm/s for a dynamic force spectrum. After each curve acquired, the glass surface was moved horizontally by at least 100 nm to expose an unused, fresh surface spot. Typically, 50000 - 100000 curves were recorded per experiment. If quantitative comparisons of absolute forces were required, a single cantilever was used to move between multiple spatially separated spots to be probed on the same surface (created using the protocol described above). To calibrate cantilevers the Inverse Optical Cantilever Sensitivity (InvOLS) was determined as the linear slope of the most probable value of typically 40 hard (>2000 pN) indentation curves. Cantilevers spring constants were calculated using the equipartition theorem method with typical spring constants between 90-160 pN nm-1. A full list of calibrated spring constants from experiments presented in this work is provided below, as the stiffness of the cantilever, may influence the complex rupture and domain unfolding forces measured. Experiments and spring constants of cantilevers for data shown:

| Measurement | Spring constant [pN/nm] | Force [pN] @ 800 nm/s |
| --- | --- | --- |
| geometry C - surf_biotin_lv_mcSAddFLN4ybbR (Figure 1D) | 56.2 | 204.2 |
| geometry N surf_biotin_lv_ybbRddFLN4mcSA2 (Figure 1D) | 120.9 | 179.9 |
| both geometries - surf_mcSA2bothmulti_lv_yFIVARbiotin (Figure 2C) | 139.2 | 187.2 / 218 |

### SMFS data analysis

Adapted from Milles et al. (18):

Data analysis was carried out in Python 2.7 (Python Software Foundation). Laser spot drift on the cantilever relative to the calibration curve was corrected via the baseline noise (determined as the last 5 % of datapoints in each curve) for all curves and smoothed with a moving median (windowsize 300 curves). The inverse optical lever sensitivity (InvOLS) for each curve was corrected relative to the InvOLS value of the calibration curve.

Raw data were transformed from photodiode and piezo voltages into physical units with the cantilever calibration values: The piezo sensitivity, the InvOLS (scaled with the drift correction) and the cantilever spring constant (k).

The last rupture peak of every curve was coarsely detected and the subsequent 15 nm of the baseline force signal were averaged and used to determine the curve baseline, that was then set to zero force. The origin of molecule extension was then set as the first and closest point to zero force. A correction for cantilever bending, to convert extension data in the position of the cantilever tip was applied. Bending was determined through the forces measured and was used on all extension datapoints (x) by correcting with their corresponding force datapoint (F) as xcorr = x - F/k.

To detect unfolding or unbinding peaks, data were denoised with Total Variation Denoising (TVD, denoised data is not shown in plots), and rupture events detected as significant drops in force relative to the baseline noise.

Rupture force histograms for the respective peaks and dynamic force spectra were assembled from all curves showing the fingerprint unfolding, or (if applicable) a specific fingerprint domain, and/or a clean complex rupture event. The most probable loading rate of all complex rupture or domain unfolding events was determined with a KDE, bandwidth chosen through the Silverman estimator. This value was used to fit the unfolding or rupture force histograms with the Bell-Evans model for each pulling velocity. A final fit was performed through the most probable rupture forces and loading rates for each pulling velocity to determine the distance to the transition state Δx0 and natural off-rate at zero force koff,0.

### Fluorescence Anisotropy Measurement

For fluorescence anisotropy measurements, biotinylated fluorescently labeled single-stranded DNA was mixed with the mcSA2 constructs in a 1:1 ratio. The change in anisotropy upon the addition of a more than 100-fold excess of biotin was recorded for 2,5 h.

Fluorescence anisotropy measurements were carried out in Corning 384 well plates. For passivation, the wells were incubated with 5 mg/ml bovine serum albumin dissolved in phosphate buffered saline (PBS) (Sigma-Aldrich, Saint Louis, USA) for 2 h. After removing the passivation solution by turning the plates upside down, the wells were flushed twice with ultrapure water.

The protein constructs were filtered with a 0.45 µm centrifuge filter (Merck Millipore, Darmstadt, Germany) according to the manufacturer’s instructions. To match the buffers, we employed Zeba Spin Desalting Columns (Thermo Scientific, Rockford, USA) with 7K MWCO using PBS to equilibrate the columns following the manufacturer’s protocol.

The concentrations of the constructs were determined with a NanoDrop 1000 (Thermo Scientific, Rockford, USA) UV-Vis spectrophotometer using the absorption peak at 280 nm and an extinction coefficient of 41035 M^-1^cm^-1^ calculated from the protein sequence using the “ExPASy: SIB bioinformatics resource portal” (21). We used 17 bp long single-stranded DNA oligonucleotides labeled with Biotin at the 5’-end and a ATTO 647N dye ot the 3’-end purchased from IBA (IBA GmbH, Göttingen, Germany).

We prepared 40 µl of 30 nM biotinylated fluorescently labeled DNA and the same amount of protein construct dissolved in PBS containing 1 mM DTT. As G-factor and measurement blank, we used 40 µl PBS with 1 mM DTT added. G-factor reference also contained 30 nM of the biotinylated fluorescently labeled DNA. After measuring the anisotropy in absence free biotin, we added 10 µl 818 µM Biotin dissolved in PBS to all wells and recorded the anisotropy every five seconds for 2.5 h.

## Acknowledgements

Support for this work was provided by the ERC Advanced Grant CelluFuel. The authors thank D.A. Pippig, F. Baumann, M.A. Jobst for helpful discussions, M. Freitag for experimental assistance, K. Erlich for proof reading and A. Kardinal and T. Nicolaus for laboratory support.

## Author contributions

M.S.B.: Conceptualization, Data curation, Software, Formal analysis, Investigation, Visualization, Writing—original draft, Writing—review and editing

L.F.M.: Conceptualization, Data curation, Software, Investigation, Visualization, Writing— original draft, Writing—review and editing

S.M.S.: Data curation, Formal analysis, Investigation, Writing—original draft, Writing—review and editing

H.E.G.: Conceptualization, Supervision, Funding acquisition, Investigation, Project administration, Writing—review and editing

